# Stable Potassium Isotopes (^41^K/^39^K) Track Transcellular and Paracellular Potassium Transport in Biological Systems

**DOI:** 10.1101/2022.01.31.478485

**Authors:** John A. Higgins, Danielle Santiago Ramos, Stefania Gili, Cornelia Spetea, Scott Kanoski, Darren Ha, Alicia A. McDonough, Jang H. Youn

## Abstract

As the most abundant cation in archaeal, bacterial, and eukaryotic cells, potassium (K^+^) is an essential element for life. While much is known about the machinery of transcellular and paracellular K transport – channels, pumps, co-transporters, and tight-junction proteins - many quantitative aspects of K homeostasis in biological systems remain poorly constrained. Here we present measurements of the stable isotope ratios of potassium (^41^K/^39^K) in three biological systems (algae, fish, and mammals). When considered in the context of our current understanding of potential mechanisms of K isotope fractionation and K^+^ transport in these biological systems, our results provide evidence that the fractionation of K isotopes depends on transport pathway and transmembrane transport machinery. Specifically, we find that passive transport of K^+^ down its electrochemical potential through channels and pores in tight-junctions at favors ^39^K, a result which we attribute to a kinetic isotope effect associated with dehydration and/or size selectivity at the channel/pore entrance. In contrast, we find that transport of K^+^ against its electrochemical gradient via pumps and co-transporters is associated with less/no isotopic fractionation, a result that we attribute to small equilibrium isotope effects that are expressed in pumps/co-transporters due to their slower turnover rate and the relatively long residence time of K^+^ in the ion pocket. These results indicate that stable K isotopes may be able to provide quantitative constraints on transporter-specific K^+^ fluxes (e.g. the fraction of K efflux from a tissue by channels vs. co-transporters) and how these fluxes change under different physiological states. In addition, precise determination of K isotope effects associated with K^+^ transport through channels, pumps, and co-transporters may provide unique constraints on the mechanisms of K transport that could be tested with steered molecular dynamic simulations.

## 1. Introduction

Archaeal, bacterial, and eukaryotic cells all concentrate potassium (K^+^). In fact, the ubiquity of elevated intracellular K^+^ has been hypothesized to provide insight into the environments (high K/Na ratios) where the first cells emerged (Mulkidjanian et al., 2012). In bacteria, intracellular K^+^ is at >100 mM by the active pumping of K^+^ from extracellular fluid (ECF) into the intracellular fluid (ICF) by transporters that use ATP or the proton motive force (Stautz et al., 2021). In eukaryotic cells, intracellular K^+^ is maintained at similarly elevated levels by the active pumping of K^+^ by plasma membrane Na,K-ATP_ase_, an energetically intensive process which can account for ~20% of basal metabolism in mammals (Rolfe and Brown, 1997). In all cells, the active transport of K^+^ into the cell establishes a steep transmembrane gradient that is a key determinant of membrane potential and a source of energy to drive action potentials, control muscle contractility, maintain cell turgor, contribute to pH homeostasis, and power ion transporters (Stautz et al., 2021; Youn and McDonough, 2009).

The regulation of K^+^ at both the cellular and organismal level, K^+^ homeostasis, is ultimately accomplished at the molecular level by transcellular and paracellular transporters (including ion pumps, channels, co-transporters) that move K^+^ across membranes and between cells (Agarwal et al., 1994; Furukawa et al., 2012; Gierth and Mäser, 2007; Youn and McDonough, 2009). Although life has evolved a diversity of molecular machines capable of K^+^ transport across cell membranes, certain aspects of cellular K^+^ transport are conserved across the major domains. For example, all known K^+^ channels are members of a single protein family whose amino acid sequence contains a highly conserved segment (the K^+^ channel signature sequence (MacKinnon, 2003)). The shared molecular machinery of K^+^ transport in biological systems reflects the fundamental role of the electrochemical potential generated by biologically maintained gradients of K^+^ across cell membranes. K^+^ pumps establish and maintain concentration gradients across cell membranes by moving K^+^ (from ECF-ICF) and Na or H (from ICF to ECF) “uphill” against their electrochemical potentials coupled to and driven by the hydrolysis of ATP. K^+^ co-transporters couple uphill K^+^ transport to “downhill” transport of another ion (e.g., Na^+^ or Cl^-^). Finally, K^+^ channels and the pores in tight-junction proteins provide an energetically favorable pathway for rapid, yet highly selective, transport of K^+^ down its electrochemical potential.

While the identity and structure of the molecular machines that maintain K^+^ homeostasis in biological systems are well-known, many quantitative aspects of K^+^ homeostasis and the specific mechanisms of K^+^ transport by pumps, channels, co-transporters, and through tight-junction proteins are unconstrained or debated. For example, structural, functional and computational studies of K^+^ channels produce conflicting results as to whether the molecular mechanism of K^+^ permeation is water mediated or occurs by direct knock-on of cations (Mironenko et al., 2021). In addition, while studies of gene expression in cells and tissues can be used to identify which K transporters are active, quantitative constraints on the fluxes of K^+^ through the different types of transporters *in vivo* are non-existent.

In natural systems potassium is made up of two stable (potassium-39 and potassium-41) and one radioactive (potassium-40) isotope. The two stable isotopes of K, ^39^K and ^41^K, constitute 93.258% and 6.730% of the total, respectively, resulting in a ratio of ^41^K/^39^K in nature of ~0.07217. Recent advances in inductively coupled plasma mass spectrometry (ICP-MS) now permit the precise quantification of deviations from the terrestrial ratio resulting from the biogeochemical cycling of potassium in nature with a precision of 1 part in 10,000 (Morgan et al., 2018; Wang and Jacobsen, 2016). Here we apply this analytical tool to study K^+^ homeostasis in three biological systems – aquatic green alga *Chlamydomonas reinhardtii* (*C. reinhardtii*), a suite of marine fish that include species that can tolerate a wide range in salinities (euryhaline) as well as those that have a restricted salinity range (stenohaline), and the terrestrial mammal *Rattus norvegicus* (*R. norvegicus*). The results in each system are interpreted as reflecting K^+^ homeostasis under normal (optimal) growth conditions. The data are presented using standard delta notation in parts per thousand (‰).

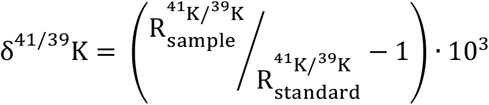

where 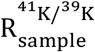 is the ratio of ^41^K/^39^K in a sample and 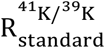 is the ^41^K/^39^K of the standard). For plants the standard is the ^41^K/^39^K of the growth medium, whereas for marine teleosts it is the ^41^K/^39^K of seawater. For *R. norvegicus*, we report the data in one of two ways. To explore total body K^+^ homeostasis we report the ^41^K/^39^K of K excretory losses normalized to the ^41^K/^39^K of the diet whereas to examine the partitioning of K isotopes between extracellular and intracellular compartments we report the ^41^K/^39^K of tissues relative to the ^41^K/^39^K of blood plasma or cerebrospinal fluid (CSF). K isotope fractionation factors are calculated as ratios of isotope ratios (e.g. α_medium/cell_ = [^41^K/^39^K_medium_] / [^41^K/^39^K_cell_]) and reported as ε values (in ‰) where S_medium/cell_ = α_medium/cell_ – 1.

## 2. Materials and Methods

### C. reinhardtii cultures

The CMJ030 wild type strain was obtained from the Chlamydomonas culture collection www.chlamycollection.org). Tris Phosphate (TP) medium was prepared according to: Gorman, D.S., and R.P. Levine (1965) Proc. Natl. Acad. Sci. USA 54, 1665-1669. The culture at an initial density of 0.5 x10^5^ cells mL^-1^ was grown in TP under continuous illumination (100 μmol photons m^-2^ s^-1^) and shaking for four days. Samples (in triplicate) containing 2 x10^7^ cells (~800 mg) were harvested and washed twice in 5 mM HEPES and 2 mM EDTA before collection and air-drying of the cell pellet. Pelletized cells (~30mg) were digested in screw-capped teflon vials on a hot plate at elevated temperatures (~75 °C) using a 5:2 mixture of HNO_3_ (68-70 vol.%) and H_2_O_2_ (30 vol.%).

### Euryhaline and stenohaline marine teleosts

Samples of teleost muscle tissue were sourced from fish markets (Nassau Seafood and Trader Joe’s in Princeton, NJ and the Fulton Fish Market in Brooklyn, NY) and research cruises (NOAA NEFSC Bottom Trawl Survey, Fall 2015 and Spring 2015). All teleosts were caught in seawater which has a uniform δ^41^K value of +0.12‰ relative to SRM3141a (Ramos et al., 2020). Samples of white dorsal muscle (100 to 3000 mg) were digested on a hot plate at elevated temperatures (~75 °C) or in a high-pressure microwave system (MARS 6) using HNO_3_ (68-70 vol.%) H_2_O_2_ (30 vol.%) in a ratio of 5:2. Major/minor element analyses for digested samples were carried out at Princeton University using a quadrupole inductively coupled plasma mass spectrometer (Thermo Scientific iCap Q). Concentrations and elemental ratios were determined using externally calibrated standards and average uncertainties (element/element) are ~10%.

### R. norvegicus experiments

All rat experiments were approved by the Institutional Animal Care and Use Committees of the University of Southern California. Two series were conducted. Series #1: Male Wistar rats (n=3, 250-275g body weight, Envigo, Indianapolis, IN) were housed in a climate controlled (22-24°C) environment with a 12 hr: 12hr light/dark cycle, and fed casein based normal K^+^ diet TD.08267 (Envigo, Indianapolis, IN) and water ad libitum for 11 days. At day 8, rats were placed overnight into metabolic cages (Techniplast, Buguggiate, Italy) with food and water ad libitum for 16-hour collection of urine and feces. On termination day (1:30-3:30PM), rats were anesthetized with an intramuscular (IM) injection of ketamine (80 mg/kg, Phoenix Pharmaceuticals, St. Joseph, MO) and xylazine (8 mg/kg, Lloyd Laboratories, Shenandoah, IA) in a 1:1 ratio. Through a midline incision, the liver, kidneys, heart, fat pads, and stomach (flushed of contents) were removed; blood was collected via cardiac puncture, spun down to separate plasma from RBCs. Then gastrocnemius, soleus, TA, and EDL skeletal muscles were dissected. All tissues were washed in ice-cold TBS to remove excess blood, weighed and snap frozen in liquid nitrogen. Series #2: Male Sprague Dawley rats (n=4, 250-300g, Envigo, Indianapolis, IN) were housed in a climate controlled (22-24°C) environment with a 12 hr: 12hr light/dark cycle and fed grain-based vivarium chow (LabDiet 5001, labdiet.com). CSF extraction procedures are as reported previously (Noble et al., 2018). In brief, Rats were deeply anesthetized using a cocktail of ketamine 90mg/kg, xylazine, 2.8 mg/kg, and acepromazine 0.72 mg/kg by intramuscular injection. A needle was lowered to below the caudal end of the occipital skull and the syringe plunger pulled back slowly, allowing the clear CSF to flow into the syringe. After extracting ~100-200 μl of CSF, the needle was raised quickly (to prevent suction of blood while coming out of the cisterna magna) and the CSF dispensed into a microfuge tube and immediately frozen in dry ice and then stored at −80 °C until time of analysis. Following CSF extraction and decapitation, whole brains with 10-15mm spinal cord extension were rapidly removed and immediately flash frozen and stored in −80°C until dissection into spinal cord, cerebrum, and cerebellum for subsequent digestion and K isotopic analysis.

### Ion chromatography and isotope ratio mass-spectrometry

K was purified for isotopic analyses using an automated high-pressure ion chromatography (IC) system. The IC methods utilized here followed those previously described in (Morgan et al., 2018; Ramos et al., 2018). The accuracy of our chromatographic methods was verified by purifying and analyzing external standards (SRM3141a and SRM70b) alongside unknown samples. Purified aliquots of K were analyzed in 2% HNO_3_ for their isotopic compositions on a Thermo Scientific Neptune Plus multi-collector inductively coupled plasma mass spectrometer (MC-ICP-MS) at Princeton University, using previously published methods (Morgan et al., 2018; Ramos et al., 2020). The external reproducibility of our protocols (chromatography and mass spectrometry) was determined through replicate measurements of international standards. Measured values of SRM70b, reported relative to SRM3141 (δ^41^K_SRM3141_) are −0.57 ± 0.17‰ (2σ; N = 59), indistinguishable from published values (Ramos et al., 2018; Ramos et al., 2020; Wang and Jacobsen, 2016). For samples analyzed once (chromatography and mass spectrometry), reported errors are the 2σ uncertainties of the external standard. In cases where samples were analyzed multiple times, reported errors in are twice the standard error of the mean (2 S.E. or 95% confidence).

## 3. Results

Results for the freshwater algae *C. reinhardtii* are shown in Figure 1. Measured δ^41^K values of the whole cells are 1.2±0.07‰ lower than the δ^41^K value of the growth medium (0‰ by definition) (95% confidence; *P* = 4.5×10^-5^).

**Figure 1.**
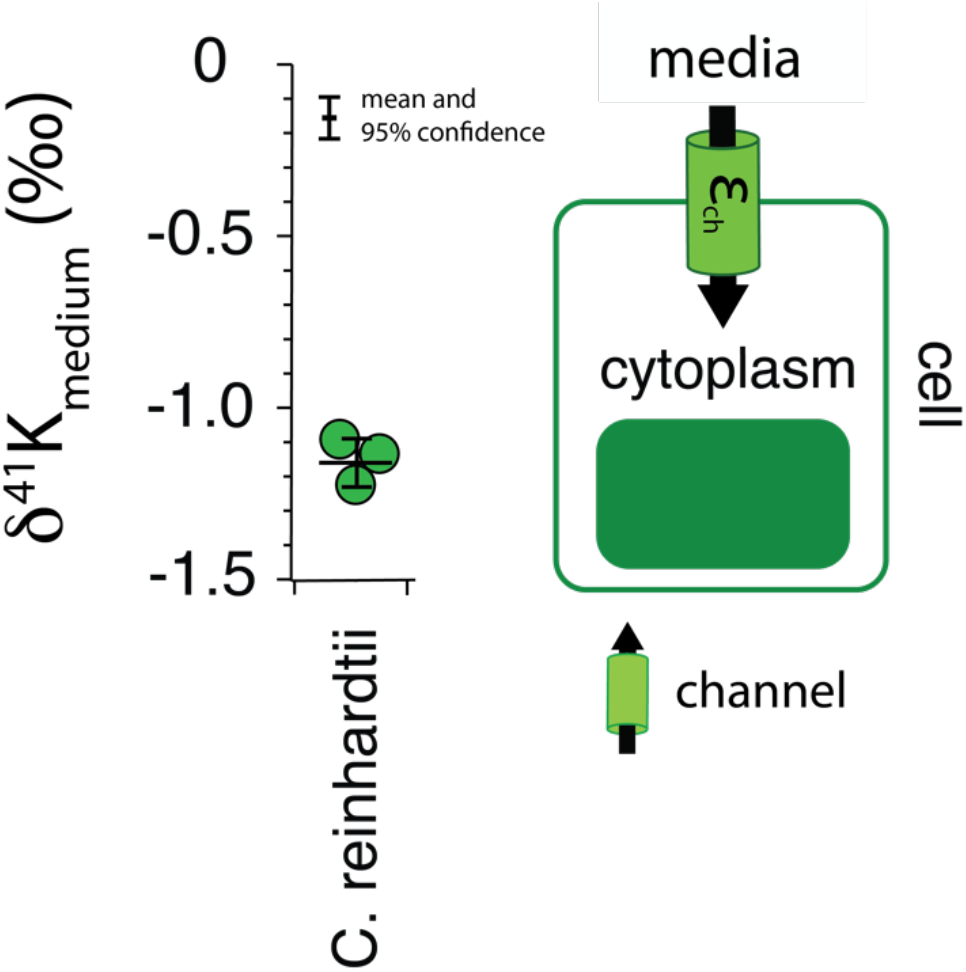
The difference in δ^41^K values between external and intracellular K^+^ in growth experiments of C. *reinhardtii* and a diagram of K^+^ transport in singlecelled algae grown under optimal conditions (See Methods) where K uptake occurs via channels that fractionate K isotopes (ε_ch_) by ~1,2‰. *P =* 4.5×1 O’^5^ by one-way ANOVA in Matlab.

Results for *R. norvegicus* are shown in Figures 2–4. Measured δ^41^K values of both urine and feces, normalized to the δ^41^K value of the rat diet (δ^41^K_diet_), indicate preferential net uptake of ^39^K relative to ^41^K across the gut epithelium leading to a positive δ^41^K_diet_ value in feces (+0.19±0.09‰, 95% confidence; *P* = 0.037; Fig. 2) and slightly negative δ^41^K_diet_ value in urine (−0.09±0.10‰, 95% confidence; *P* = 0.1895).

**Figure 2.**
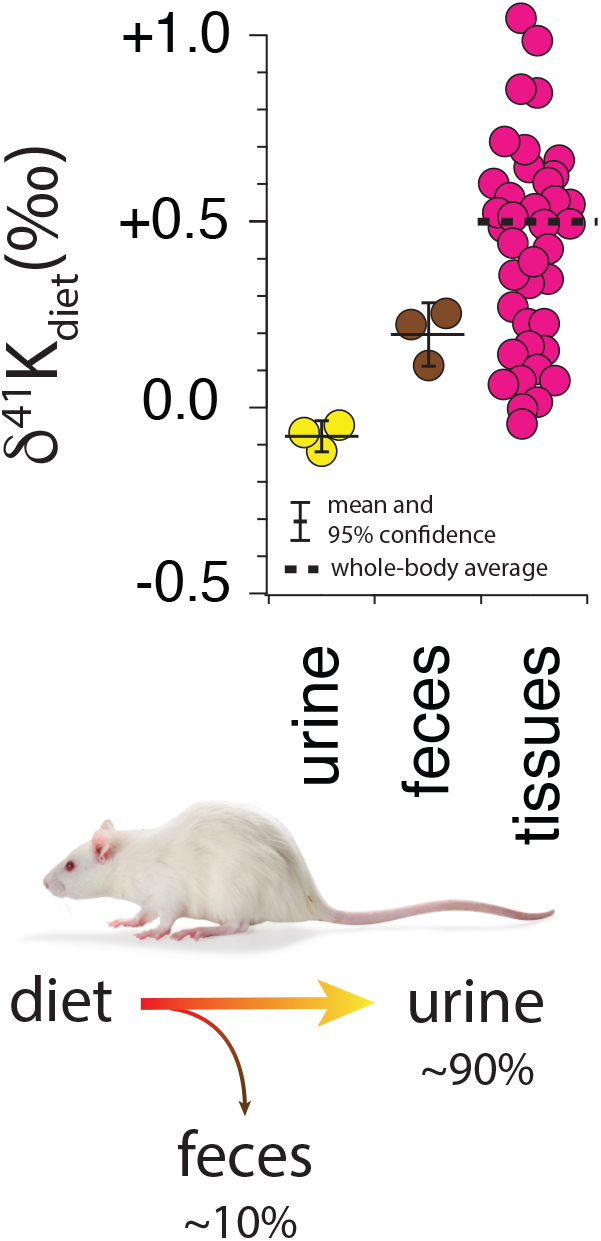
Whole-body K isotope mass balance in *R. norvegicus* on a controlled diet. Measured δ^41^K values of urine (*P* = 0.19) and feces (*P* = 0.037) relative to diet indicate a preferential uptake of ^39^K in the gut. The total range in δ^41^K_diet_ values in the tissues of R. *norvegicus* is ~1‰ with an estimated whole-body average δ^41^K_diet_ value of approx. +0.5‰, consistent with preferential loss of ^39^K in urine.

**Figure 3.**
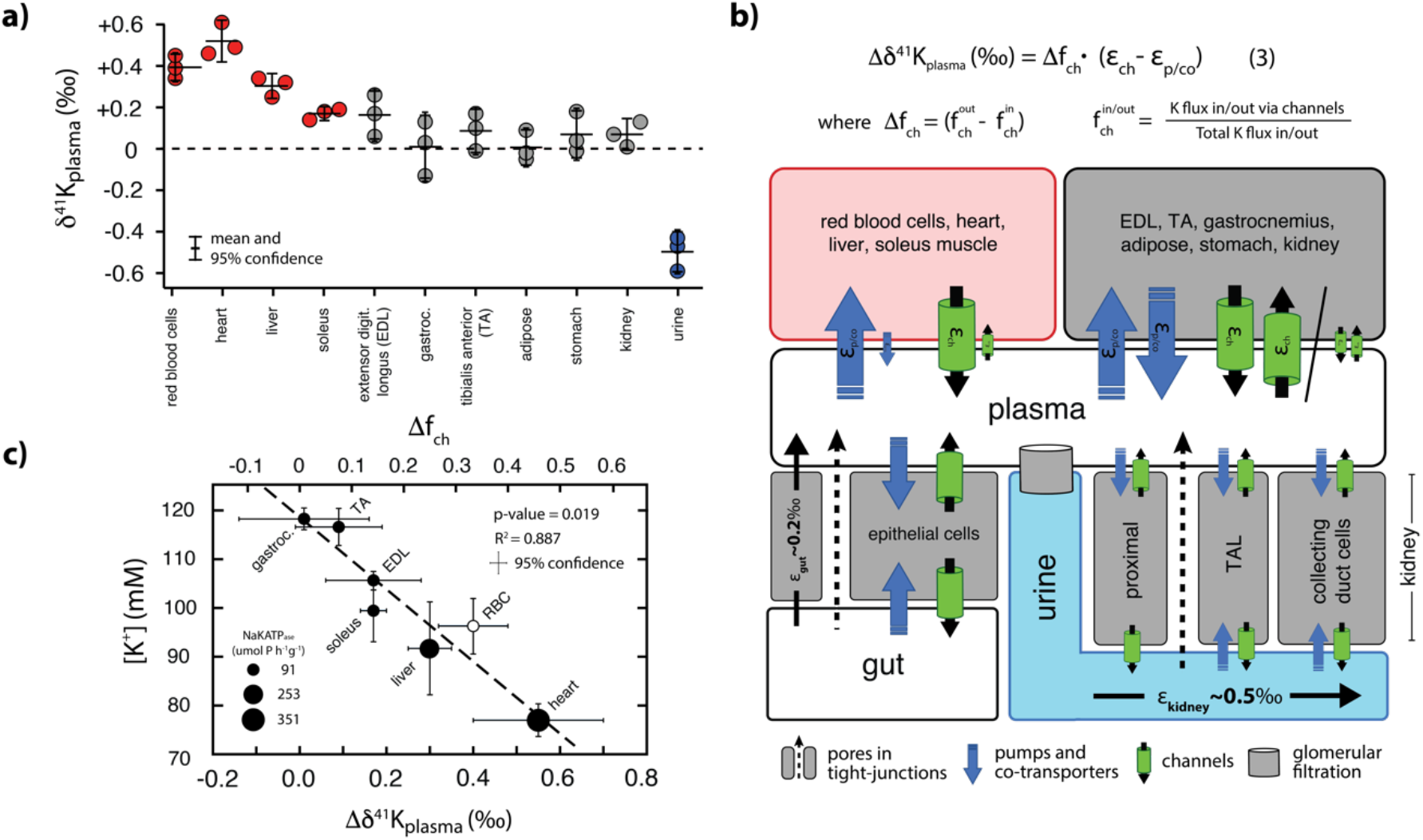
a) K isotopic composition of tissues/fluids *Rattus norvegicus* normalized to plasma (δ^41^K_plasma_ = 0‰). δ^41^K_plasma_ values that are positive 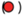 indicate that net K^+^ transport out of the tissue/cell/fluid is enriched in ^39^K relative to K^+^ transport into the tissue/cell/fluid. δ^41^K_plasma_ values that are indistinguishable from 0 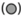 indicate that the K isotope fractionation associated with K^+^ transport into and out of the cell/tissue/fluid are either 0 or equal in magnitude (and thus cancel). δ^41^K_plasma_ values that are negative 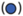 indicate that net K^+^ transport into the cell/tissue/fluid is enriched in ^39^K relative to K^+^ transport out of the cell/tissue/fluid. **b)** A diagram of K isotope mass balance including transport by (1) K-channels, (2) K-pumps/co-transporters, (3) through pores in tight-junctions, and (4) glomerular filtration. Equation (3) solves for the δ^41^K value of a tissue/cell relative to blood plasma assuming steady-state K isotope mass balance with bi-directional K^+^ transport by both channels and pumps/co-transporters. Variables include 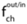, the fraction of total K^+^ transport into/out of the tissue/cell that occurs through K^+^ channels, Δf_ch_, the difference in the fraction of K transport via channels 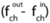, ε_ch_, K isotope fractionation associated with K channels, and ε_p/co_, K isotope fractionation associated with pumps/co-transporters. K isotope fractionation associated with K^+^ uptake in the gut (ε_gut_) and K^+^oss in urine (ε_kidney_) reflect the flux-weighted average of both paracellular and transcelluar K^+^ transport. For example, K isotope fractionation in the kidney will be associated with both re-absorption and secretion of K^+^ in the proximal tubule, the thick ascending limb (TAL), and cells in the collecting ducts of the kidney, c) Correlation between intercellular [K^+^] and Δδ^41^K_plasma_ values for a subset of tissues from *R. norvegicus*. Also shown are the measured activities of NaKATPase (μmol P h ‘ g ‘) from (49) and values of Δf_ch_ from eqn (3).

**Figure 4.**
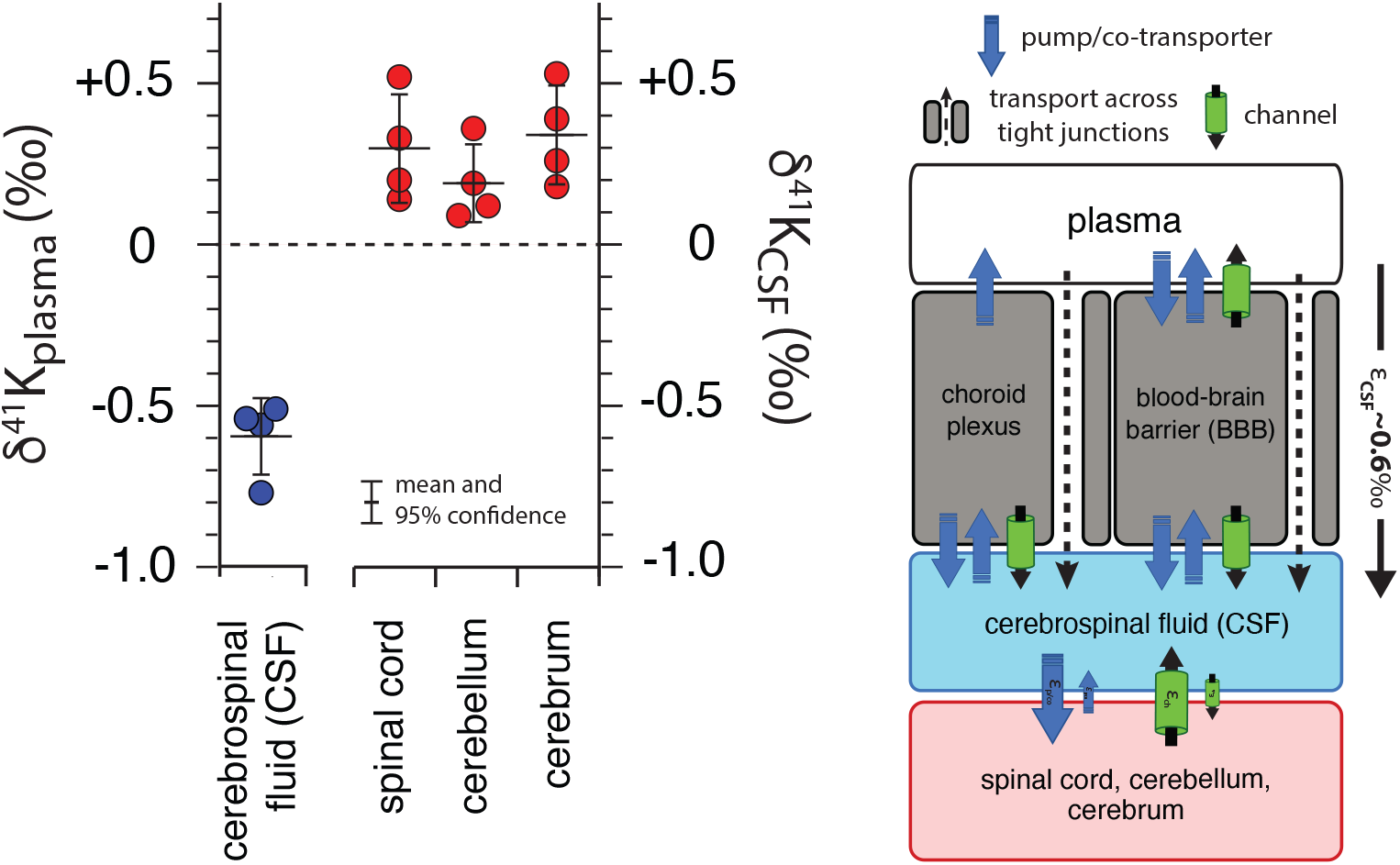
K isotopic composition of cerebrospinal fluid from *R. norvegicus* raised on a controlled diet normalized to blood plasma (δ^41^K_plasma_ = 0‰) and brain tissues normalized to cerebrospinal fluid (δ^41^K_CSF_ = 0‰) together with a diagram of K homeostasis across the choroid plexus and blood brain barrier (BBB) from (50) including transport by (1) K-channels, (2) K-pumps/co-transporters, and (3) transcellularly through pores in tight-junctions. δ^41^K_CSF_ values that are positive 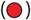 indicate that net K transport out of brain tissues (spinal cord, cerebellum, and cerebrum) and into CSF is enriched in ^39^K relative to K transport from CSF into these tissues. The negative δ^41^K_plasma_ values 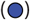 for CSF indicate that net K transport into CSF from blood plasma is enriched in ^39^K relative to K transport from CSF to blood plasma andreflects the flux-weighted average of the tranporters that dominate K exchange across both the choroid plexus and BBB.

Measurements of K isotopes of various tissues in *R. norvegicus* (except brain tissues) are normalized to the δ^41^K value of the blood plasma (δ^41^K_plasma_), as this represents the extracellular K concentration bathing the tissues. For brain tissues, measured δ^41^K values are normalized to CSF (δ^41^K_CSF_) for the same reason. The overall range in δ^41^K_plasma_ values between different tissues and fluids is ~1‰ (Figs. 3,4); δ^41^K values of the following tissues are elevated in ^41^K relative to blood plasma (positive δ^41^K_plasma_ values): red blood cells (+0.40±0.08‰, *P* = 0.005), heart (+0.55±0.15‰, *P* = 0.0047), liver (+0.30±0.05‰, *P* = 0.01), and soleus muscle (+0.17±0.03‰, *P* = 0.05). Similarly, brain tissues are elevated in ^41^K relative to CSF (positive δ^41^K_CSF_ values): cerebrum (+0.34±0.15‰, *P* = 0.012), spinal cord (+0.30±0.17‰, *P* = 0.026), and cerebellum (+ 0.21±0.11‰, *P* = 0.037). In contrast, urine (−0.50±0.10‰, *P* = 0.0019) and CSF (−0.59±0.12‰, *P* = 6.2×10^-5^) are characterized by negative δ^41^K_plasma_ values. In another category are tissues with δ^41^K_plasma_ values that are statistically indistinguishable from zero including: kidneys (+0.07±0.07‰, *P* = 0.31), adipose tissue (+0.01±0.08‰, *P* = 0.92), extensor digitorum longus (EDL, +0.17±0.11‰, *P* = 0.099) gastrocnemius (+0.01±0.15‰, *P* = 0.92), and tibialis anterior (TA, +0.09±0.1‰, *P* = 0.30).

Results for white muscle tissue from a suite of stenohaline and euryhaline marine fish, reported relative to the ^41^K/^39^K of seawater (δ^41^K_seawater_ = 0‰) are shown in Figure 5a. The total range in muscle δ^41^K values is ~2‰ (+1‰ to −1‰). Stenohaline species including *Gadus morhua* (Atlantic Cod), *Peprilus striacanthus* (Butterfish), Xiphias gladius (Swordfish), *Pseudopleuronectes americanus* (Winter Flounder), and *Hippoglossus stenolepis* (Pacific Halibut) are characterized by δ^41^K_seawater_ values that are uniformly negative whereas the euryhaline species *Oncorhynchus kisutch* (Coho Salmon), *Oncorhynchus tshawytscha* (King Salmon), and *Oncorhynchus nerka* (Sockeye Salmon) are characterized by δ^41^K_seawater_ values that are close to zero or positive. When species are grouped by salinity tolerance, average measured δ^41^K_seawater_ values of the two groups are −0.58±0.09‰, and 0.29±0.18‰ for stenohaline and euryhaline species, respectively (95+ confidence; *P* = 1.4×10^-10^).

**Figure 5.**
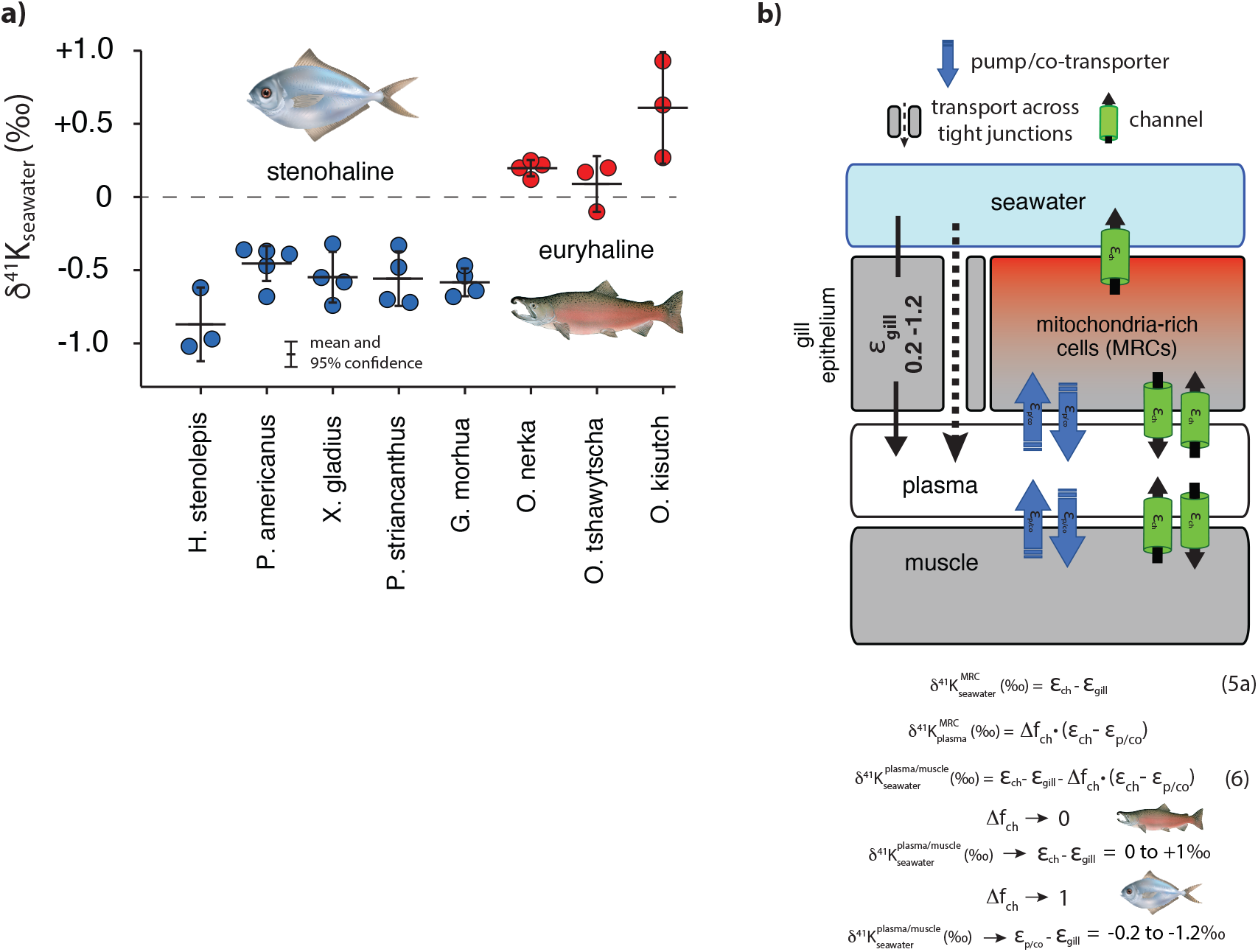
**a)** The difference in δ^41^K values between seawater and dorsal white muscle of various stenohaline and euryhaline marine fish. Muscle [K^+^] and δ^41^K_seawater_ values are listed in Table S1. *P* = 1.4×10^-10^ for the difference in δ^41^K_seawater_ value between stenohaline and euryhaline teleosts. **b)** A model of of K isotope mass balance in a marine fish. K^+^ is supplied across the gill epithelium via transport through pores in tight-junction proteins and lost from mitochondria-rich cells (MRCs) through channels (equation 5). K^+^ cycling within MRCs is modeled using eqn. (3). Combining eqn. (3) and eqn. (5) yields eqn. (6), the δ^41^K value of plasma/muscle relative to seawater as a funciton of ε_ch_, ε_gill_, Δf_ch_ of MRC’s and ε_p/co_. As Δf_ch_ → 0, δ^41^K of fish muscle → ε_ch_ - ε_gill_ (≥ 0‰, euryhaline), whereas as Δf_ch_ → 1, δ^41^K of fish muscle → ε_p/co_ - ε_gill_ (< 0‰, stenohaline).

## 4. Discussion

The shared machinery of K^+^ transport across the domains of life suggests that the variability in K isotopes observed in biological systems can be reduced to a consideration of 1) how different classes of K^+^ transporters - tight junctions, channels, pumps, and co-transporters - discriminate between ^39^K and ^41^K and 2) how these transporters are assembled into a homeostatic system. The extent to which K^+^ transporters fractionate K isotopes will, in turn, depend on the mechanics and selectivity of the transporter and whether the binding of dissolved K^+^ to transporter is the rate-limiting step. In particular, K^+^ channels and pores in tight-junction proteins are both associated with selective and rapid (near the limit of diffusion) transport of K^+^ where the rate-limiting step is the binding of dissolved K^+^ to the transporter/selectivity filter. These conditions favor kinetic isotope effects which may arise from either partial or full desolvation of K^+^ as it binds to the selectivity filter (Berneche and Roux, 2003; Hofmann et al., 2012; Kopec et al., 2018; Yu et al., 2010), differences in ionic radius of ^39^K and ^41^K (size selectivity (Christensen et al., 2018)), or some combination of the two. Critically, the magnitude of the kinetic isotope effects associated with both desolvation and/or size selectivity appear to be large, with estimates from laboratory experiments and molecular dynamic simulations ranging from ~1 to 2.7‰ (Hofmann et al., 2012).

In contrast to channels and pores in tight-junction proteins, transport of K^+^ by pumps and co-transporters is relatively slow (per unit transporter, (Roux, 2017)), requires the simultaneous binding of multiple ions (e.g. 2 K^+^ ions in the case of Na,K-ATP_ase_), and is associated with mechanisms of self-correction that prevent the pump cycle from proceeding if incorrect ions bind to the ion pocket (Rui et al., 2016). As a result, although the binding of K^+^ to the active site of a pump or cotransporter may also involve desolvation, it is not the rate-limiting step in K^+^ transport. In this case, the isotopic composition of K^+^ that is transported by pumps and co-transporters may reflect isotopic equilibration between K ^+^ in the ion pocket and the fluid due to the longer residence time of K^+^ in the ion pocket prior to occlusion. As equilibrium K isotope effects tend to be small (Li et al., 2017; Ramos et al., 2018; Zeng et al., 2019), we hypothesize that K^+^ transport by pumps and co-transporters will be associated with less isotopic fractionation.

In the following sections we show how our results for the three studied biological systems (algae, fish, mammals) can be interpreted within a single framework where K^+^ transport by channels and pores in tight-junction proteins are associated with large (kinetic) K isotope effects whereas K^+^ transport by pumps and/or co-transporters is associated with smaller (equilibrium) K isotope effects. In simple biological systems (e.g. single celled alga) linking K isotope fractionation to the machinery of K^+^ transport is relatively straightforward due to the small number of pathways involved and the unidirectional nature of K^+^ transport. In more complex biological systems (e.g. higher plants, fish, and mammals) discerning transporter-specific fractionation of K isotopes is more difficult for at least two reasons. First, our understanding of K homeostasis at both the tissue and organism level is incomplete. Second, our measured K isotopic differences (e.g. between plasma/CSF/seawater and tissue) only provide information on net K isotope fractionation associated with bi-directional K^+^ transport at steady-state. In spite of these limitations, our results support a framework for K isotope fractionation in biological systems that can provide new quantitative insights into the mechanisms of K^+^ transport and the dynamics of K homeostasis.

### 4.1 Algae

As an essential macronutrient in all plants and the most abundant cation in the cytoplasm, K^+^ contributes to electrical neutralization of anionic groups, membrane potential and osmoregulation, photosynthesis, and the movements of stomata (Gierth and Mäser, 2007; Marchand et al., 2020). It is well-established that K^+^ channels play prominent roles in K^+^ uptake (Lebaudy et al., 2007). For example, under normal growth conditions ([K^+^]_ext_ ~1 mM; (Epstein et al., 1963; Kochian and Lucas, 1982; Malhotra and Glass, 1995)) K^+^ uptake in plants is dominated by transport via inward-rectifying K^+^ channels electrically balanced by the ATP-driven efflux of H^+^ (Britto and Kronzucker, 2008; Hirsch et al., 1998). Considered in this context our results for whole cells of *C. reinhardtii* can be interpreted as reflecting K isotope fractionation associated with uptake via K^+^ channels (ε_in_ = δ^41^K_medium_ – δ^41^K_cell_ = ~1.2‰; Figure 1). Although we cannot guarantee that K uptake occurs exclusively via K^+^ channels, this result represents our best estimate of a transporter-specific K isotope effect (ε_ch_). The magnitude of this effect is smaller than modeled estimates of the kinetic isotope effect associated with ion desolvation (Hofmann et al., 2012) and of the same order as hypothesized kinetic isotope effects associated with size-selectivity (Christensen et al., 2018).

### 4.2 Higher Plants

Our result for K isotope fractionation associated with K^+^ uptake in *C. reinhardtii* is similar in sign, though somewhat larger in magnitude, than the K isotope effect associated with K^+^ uptake estimated by Christensen et al. (2018) in experiments involving higher plants. Those authors analyzed the δ^41^K values of the roots, stems, and leaves of *Triticum aestivum* (wheat), *Glycine max* (soy) and *Oryza sativa* (rice) grown under hydroponic conditions and observed systematic differences in the δ^41^K value of the different reservoirs. In particular, roots, stems, and leaves exhibited increasingly negative δ^41^K values (Figure S1). Compared to the freshwater alga *C. reinhardtii*, quantifying transport-specific K isotope effects in higher plants is more complex as the δ^41^K value of each individual compartment (root, stem, leaf) reflects a balance between isotopic sources and sinks (e.g. the δ^41^K value of the root will depend on K isotope effects associated with both net K^+^ uptake as well as translocation and recycling). Using a model of K isotope mass balance that includes assumptions regarding plant growth and the partitioning of K^+^ fluxes between translocation and recycling, Christensen et al. (2018) estimated large K isotope effects for uptake (ε_in_ ~0.7 to 1.0‰) and translocation from root to stem (~0.6‰), but smaller isotopic effects (0 to 0.2‰) for translocation of K^+^ from stem to leaf and recycling of K^+^ from leaf to root. Interestingly, in addition to their important role in K^+^ uptake, K^+^ channels have also been shown to be involved in translocation (root to stem; (Gaymard et al., 1998)) providing a potential explanation for the relatively large K isotope effect observed by Christensen et al. (2018) for this process. In contrast, translocation and recycling of K^+^ from leaf to root, a process that also involves K^+^ channels (Lacombe et al., 2000; Pilot et al., 2001), does not appear to result in significant fractionation of K isotopes. This could be due to the increased importance of other types of K^+^ transporters in this process, a reduction in the expression of K isotope fractionation by channels due to rapid internal recycling of K^+^ between stem and leaf (compared to K^+^ transport from root to stem), or some combination of the two.

### 4.3 Terrestrial Mammals

K^+^ homeostasis in terrestrial mammals reflects the balance between K^+^ gained from diet and K^+^ lost in urine and feces (Figure 2, Figure 3a). This balance is largely achieved by the kidneys and colon, which possess a remarkable ability to sense a change in K^+^ in the diet and then appropriately adjust K^+^ loss in response. ECF (which includes blood plasma), the reservoir through which internal K^+^ is exchanged, represents only 2% of total body K^+^. Though small, this reservoir is tightly regulated to maintain membrane potential as indicated by the narrow range of normal ECF [K^+^] (~ 3.5 – 5 mEq/L,)(Boron, 2005; McDonough and Youn, 2017). Of the remaining 98% of total body K^+^, 75% resides in muscle tissues ([K^+^] ~ 130 mEq/L) and 23% in non-muscle tissues. Some tissues, particularly skeletal muscle, are critical to K^+^ homeostasis by providing a buffering reservoir of K^+^ that can take up K^+^ after a meal and altruistically donate K^+^ to ECF to maintain blood plasma levels during fasting. Cerebrospinal fluid (CSF), the body fluid that surrounds the brain and spinal cord of all vertebrates, merits special mention as its K^+^ content is even more tightly regulated than ECF (Bradbury and Davson, 1965).

#### 4.3.1. External K isotope mass balance in terrestrial mammals

Roughly ~80-90% of K in the diet is lost in urine with the remainder lost in feces (Agarwal et al., 1994). Uptake of K^+^ across gut endothelial cells occurs largely paracellularly through pores in tight-junction proteins like caudin-15 (Garcia-Hernandez et al., 2017) as a result of the transmucosal electrical potential difference (Barnaby and Edmonds, 1969). However, transcellular K^+^ uptake (via pumps) and secretion (via channels) has also been shown to occur in gut epithelial cells (Foster et al., 1984). As a result, K isotope fractionation associated with net K uptake in the gut will reflect the flux-weighted average of the K isotope fractionation associated with both transcellular and paracellular K^+^ transport. The observation that feces is characterized by a positive δ^41^Kdiet value and urine with a slightly negative δ^41^K_diet_ value (Figure 2) indicates that the net K isotope effect associated with K uptake under these conditions (ε_gut_) is ~0.2‰ (Figure 3b). Relative to the diet, internal tissues of *R. norvegicus* exhibit a 1‰ range from −0.04‰ for the cerebellum to +1.05‰ for the heart (Figure 2). Assuming an internal K^+^ distribution that is 75/15/10 muscle/adipose/other tissues, a reasonable average whole-body δ^41^K_diet_ value of *R. norvegicus* is ~+0.5‰. As net uptake of K^+^ in the gut prefers ^39^K, the elevated average whole-body δ^41^K_diet_ of *R. norvegicus* requires that there be even greater K isotope fractionation (favoring ^39^K) associated with K^+^ loss in the urine (see below).

#### 4.3.2 Internal K isotope mass balance in terrestrial mammals

As shown in Figures 3a and 4, measured δ^41^K values of the 16 different tissues and fluids analyzed in *R. norvegicus* fall into 3 distinct categories relative to ECF (δ^41^K_plasma_ or δ^41^K_CSF_): those with positive δ^41^K_plasma/CSF_ values (red blood cells, heart, liver, soleus muscle and brain tissues), those with δ^41^K_plasma/CSF_ values that are close to 0 (stomach, adipose tissue, kidney, gastrocnemius, EDL and TA muscles), and those with negative δ^41^K_plasma/CSF_ values (urine and CSF).

The timescale for K^+^ turnover in the studied tissue/cell/fluid reservoirs is rapid (e.g. 0.9-10%/min; (Jones et al., 1977; Terner et al., 1950)). As a result, for all measured tissues/cells, the δ^41^K values in Figure 3a can be interpreted as reflecting steady-state K isotope mass balance between the tissue/cell/fluid and the relevant fluid (blood plasma/ECF or CSF). At isotopic steady-state, the δ^41^K value of K^+^ entering the reservoir must equal to δ^41^K value of K^+^ leaving the reservoir. As a result, reservoirs with δ^41^K_plasma/CSF_ values that differ from 0‰ require that net K^+^ transport in one direction results in greater fractionation of K isotopes than net K^+^ transport in the opposite direction. For example, reservoirs with positive δ^41^K_plasma_ values including red blood cells, heart, liver, and soleus require that K^+^ transport from ICF to plasma or ECF is associated with a larger K isotope effect than K^+^ transport from ECF to ICF. The same relationship is observed between brain tissues and CSF (Figure 4). Similarly, reservoirs with δ^41^K_plasma_ values that are close to 0‰ require that net K^+^ transport in both directions does not fractionate K isotopes, i.e., the isotope effects must be of equal magnitude and sign and cancel. These include kidney, adipose tissue, stomach, and gastrocnemius and TA muscles. Finally, the negative δ^41^K_plasma_ values for CSF requires that net K^+^ transport from ECF to CSF through the choroid plexus and blood brain barrier (BBB) is characterized by a K isotope effect that is ~0.6‰ greater than the K isotope effect associated K^+^ transport from CSF to ECF (Figure 4).

Quantitatively linking the observed isotopic differences between ECF and ICF of various tissues/cells to the machinery of K^+^ homeostasis in complex biological systems is complicated by both an incomplete understanding of the machinery involved and limitations of steady-state isotopic mass balance. In particular, although much is known about the identity and molecular structure of the machinery that maintains K^+^ homeostasis, quantitative information on how each transporter contributes to the gross fluxes of K^+^ between ICF and ECF at the resting membrane potential is lacking. For example, in red blood cells, probably the best understood with regard to the machinery of K^+^ homeostasis due to the fundamental role it plays in the regulation of cell volume and longevity (Tosteson and Hoffman, 1960), elevated intercellular K^+^ is believed to be maintained by a ‘pump-leak’ mechanism where the pump is Na,K-ATP_ase_ and the leak is K^+^ channels (Gardos, 1958). However, red blood cells also show activity for co-transporters such as NKCC (Duhm, 1987; Sachs, 1971) and KCC (Franco et al., 2013), as well as inward-rectifying K^+^ channels (Hibino et al., 2010), and exactly how much K^+^ is transported by which transporter at the resting membrane potential is not well known.

In order to evaluate whether or not differences in measured δ^41^K_plasma_ values reflect quantitative differences in K^+^ fluxes through various transporters, we developed a generic model of K homeostasis (Figure 3b) that assumes 1) the tissue/cell is at the resting membrane potential and K is at isotopic steady-state between ECF and ICF, 2) all channels fractionate K isotopes identically (ε_ch_), 3) all pumps and co-transporters fractionate K isotopes identically (ε_p/co_), and 4) pumps, co-transporters, and channels may transport K^+^ in either direction (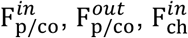 and 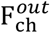):

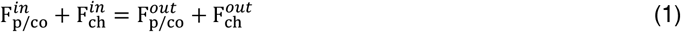

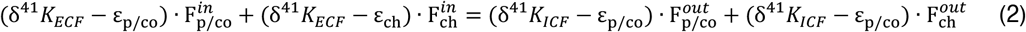

where ε_ch_, and ε_p/co_ are the K isotope fractionation factors associated with channels and pumps/co-transporters, respectively. Solving for the Δδ^41^K_plasma_ value of the ICF (δ^41^K_ICF_ - δ^41^K_ECF_) yields:

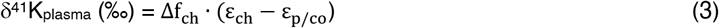

where 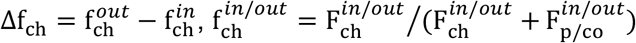. Assuming, based on our results for *C. reinhardtii*, that ε_ch_ ~ 1.2‰ and speculating, based on analogues with equilibrium K isotope effects associated with mineral precipitation in aqueous systems (Li et al., 2017; Ramos et al., 2018; Zeng et al., 2019), that ε_p/co_ ~ 0‰, reduces eqn (1) to:

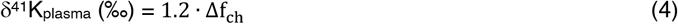

As 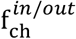 represent fractions of a flux, the value of Δf_ch_ is bounded at ±1; +1 representing a perfect ‘pump-leak’ mechanism, −1 representing a perfect ‘leak-pump’ mechanism, and 0 consistent with either a ‘pump-pump’ or ‘leak-leak’ mechanism. Accordingly, Δδ^41^K_plasma_ values are bounded at ±1.2‰, with +1.2‰ representing a perfect ‘pump-leak’ mechanism, −1.2‰ representing a perfect ‘leak-pump’ mechanism, and 0‰ representing either a ‘pump-pump’ or ‘leak-leak’ mechanism. The observation that the Δδ^41^K_plasma_ values of all internal reservoirs are ≥ 0 suggests that Δf_ch_ ≥ 0, a result that is qualitatively consistent with the idea of elevated intercellular K^+^ being maintained by a ‘pump-leak’ mechanism. However, δ^41^K_plasma_ values of the analyzed reservoirs also exhibit a considerable range, from ~0‰ for many skeletal muscles to +0.55‰ for the heart. In the context of eqn. (1), these differences can be explained by an increase in the relative importance of K^+^ loss via channels. Skeletal muscles appear to maintain K homeostasis by something closer to a ‘pump-pump’ or ‘leak-leak’ mechanism whereas red blood cells, the liver and the heart approach ~50% of a perfect ‘pump-leak’ mechanism.

Two lines of evidence provide additional independent support for our interpretation of δ^41^K_plasma_ values as quantitative indicators of the relative importance of K^+^ loss from ICF via channels (eqn. (3)). First, measured δ^41^K_plasma_ values are correlated with the activity of NaKATP_ase_ (umol P/h per g tissue) in tissues where this activity is related to the maintenance of intercellular [K^+^]. In particular, NaKATP_ase_ activity increases from 91±15 in skeletal muscles, to 253±8 in the liver, and 351±23 in the heart (Gick et al., 1993). All else held constant, this increase in NaKATP_ase_ activity (K^+^ sources to the ICF via pumps) will lower 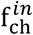 and increase Δf_ch_, in agreement with observations. Second, measured δ^41^K_plasma_ values also correlate with intercellular [K^+^] (Figure 4c). Because intercellular [K^+^] reflects the balance between K^+^ the sources and sinks to the ICF, one possible explanation for lower intercellular [K^+^] is a reduction in K^+^ supply and/or an increase in K^+^ loss. As K^+^ supply from NaKATP_ase_ increases (see above), the most likely alternative explanations are a decline in K^+^ supply from inward rectifying K^+^ channels and/or an increase in K^+^ loss via outward rectifying K^+^ channels. Both of these processes increase Δf_ch_ and result in higher δ^41^K_plasma_ values, again in agreement with observations.

##### 4.3.2.1 Urine

Unlike the internal K^+^ reservoirs discussed above, all of which are interpreted as independent homeostatic systems at isotopic steady-state with respect to ECF (plasma), the loss of K^+^ through the urine represents the end product of a series of steps each of which may contribute to the observed net K isotope fractionation (e.g. Figure 3b; ε_kidney_~0.50‰). These are 1) glomerular filtration, 2) reabsorption along the proximal tubule and the thick ascending limb of the loop of Henle, and 3) secretion and reabsorption by principal and intercalated collecting duct cells. Although a detailed description of the potential K isotope effects associated with each step is beyond the scope of this manuscript, a brief description follows. Glomerular filtration is not expected to fractionate K isotopes as the slit diaphragms freely filter ions and small molecules. A large fraction (~85%) of the filtered K^+^ is subsequently reabsorbed via both paracellular and transcellular routes in the proximal tubule and thick ascending limb (TAL). The residual K^+^ is passed along to the collecting ducts where K^+^ is added prior to excretion as urine. The addition of K^+^ in the collecting ducts occurs transcellularly, via ROMK channels in principal cells and BK channels in intercalated cells (McDonough and Youn, 2017). Although we expect K^+^ channels to be associated with a large K isotope effect (ε_ch_ = 1.2‰), the extent to which this determines the δ^41^K_plasma_ value of urine depends on 1) the δ^41^K_plasma_ value of the residual K^+^ exiting the TAL (after reabsorption but before secretion) and 2) internal K^+^ cycling within the principal and intercalated cells (e.g. eqn (3)). The observation that ε_kidney_ is only ~ 0.50‰ suggests that either the residual K^+^ exiting the TAL has an elevated δ^41^K_plasma_ value due to preferential removal of ^39^K during reabsorption and/or the δ^41^K_plasma_ value of ICF in principal and intercalated cells elevated, for example due to high values of Δf_ch_.

##### 4.3.2.2 Cerebrospinal fluid (CSF)

K^+^ in CSF reflects a balance between paracellular and transcellular K^+^ transport across endothelial cells at the blood-brain-barrier (BBB) and paracellular K^+^ transport of across epithelial cells of the choroid plexus (Hladky and Barrand, 2016). Gross fluxes of K^+^ into the brain across the BBB are 4x larger than those associated with the choroid plexus (Katzman et al., 1965), suggesting that the observed net K isotope effects may be largely due to fractionation associated with transport across the BBB. However, K^+^ transport from ECF to CSF through the choroid plexus is thought to occur by paracellular routes through pores in tight junctions (Figure 4), a process that we expect to fractionate K isotopes and thus may contribute to the observed negative δ^41^K_plasma_ values for CSF. With regards to K^+^ transport across the BBB, both paracellular (through tight-junction pores) and transcellular (through BBB endothelial cells) routes may be important (Hladky and Barrand, 2016). Again, we expect paracellular K^+^ transport through tight junction pores to be associated with a larger K isotope effect whereas any K isotope effects associated with transcellular transport will depend on the internal cycling of K^+^ (and associated isotope effects) within endothelial BBB cells (pumps, (Betz et al., 1980); co-transporters, (Foroutan et al., 2005); and channels, (Van Renterghem et al., 1995)). Overall, the observation of large K isotope fractionation associated with the transport of K^+^ from plasma to CSF (Fig. 5; ε_CSF_ = 0.59 ± 0.12‰) requires that either 1) there is a large K isotope effect associated with transcellular K^+^ transport across endothelial BBB cells or 2) transport of K^+^ from plasma to CSF is dominated by paracellular routes in both the choroid plexus and across the BBB.

### 4.4 Stenohaline and Euryhaline Marine Fish

K^+^ homeostasis in both euryhaline and stenohaline marine fish is linked to ionic and osmotic regulation (Figure 5). While K^+^ sources include ingestion of seawater and diet (Hickman Jr, 1968), by far the largest K^+^ source is transport across the gills (Maetz, 1969), which are permeable to monovalent cations (Na^+^, K^+^) and anions (Cl^-^). Fish balance this salt intake by actively secreting Na^+^, K^+^ and Cl^-^ through mitochondria-rich cells (MRCs) of the gills and paracellularly through tight-junction proteins (claudins) in the gill epithelium (Evans et al., 2005; Kolosov et al., 2013). K^+^ loss is not well-understood but recent discoveries of apical ROMK channels indicate that transcellular secretion through MRCs is likely an important pathway (Furukawa et al., 2012). Both stenohaline and euryhaline marine fish possess this capability but euryhaline marine fish have evolved the ability to adapt this machinery to a wide range of water salinities by adjusting expression of ROMK and tight junction claudins (Furukawa et al., 2012; Furukawa et al., 2014; Furukawa et al., 2015; Kolosov et al., 2013).

In seawater adapted stenohaline and euryhaline marine fish, K^+^ homeostasis can be approximated as a balance between gain of K^+^ across tight-junctions in the gill epithelium and loss of K^+^ through apical ROMK channels in MRCs (Figure 5b). Other potential sources and sinks of K^+^ including the ingestion of seawater, diet, and excretion are either small compared to the fluxes of K^+^ across the gills (Maetz, 1969) or transient in nature and unlikely to explain the systematic difference we observe between the δ^41^K_seawater_ values of stenohaline and euryhaline fish. Furthermore, as most of the total K^+^ content of fish resides in muscle tissue, the δ^41^K_seawater_ value of the muscle can be used as a reasonable approximation of the δ^41^K_seawater_ value of the whole organism (Figure 5b). At steady-state 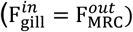, the δ^41^K_seawater_ value of the whole organism will reflect the balance between K isotope fractionation associated with K^+^ sources (paracellular transport across the gills, ε_gill_) and K^+^ sinks (transcellular transport through ROMK channels in MRCs, ε_ch_):

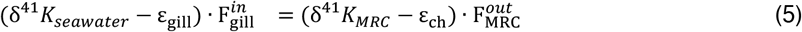

which reduces to:

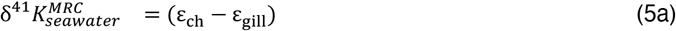

where 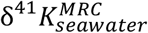 is the difference between the δ^41^K value of the MRCs and seawater. The δ^41^K value of MRCs can be related to the δ^41^K value of the whole fish by combining equation (5a) with equation (3) and assuming 1) that the flux of K^+^ out of MRCs via apical ROMK channels is small compared to the gross fluxes of K^+^ into/out of MRCs associated with K homeostasis and 2) the δ^41^K value of plasma is similar to muscle (a reasonable assumption for most muscles in *R. norvegicus*, Figure 3b). In this case:

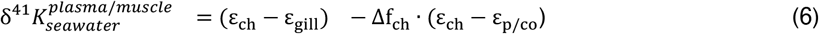

where Δf_ch_ is the relative difference in K loss via channels in MRCs. Although we do not independently know the value of ε_gill_ and suspect that it may vary systematically between stenohaline and euryhaline marine fish, we speculate that ~0.2 to 1.2‰ represents a plausible range based on results for epithelial cells in the gut of *R. norvegicus* (Figure 3b) and the maximum K isotope fractionation expected for channels (ε_ch_). Values of ε_ch_ and ε_p/co_ are assumed to be 1.2‰, and 0‰, respectively. Given these values, inspection of equation (6) indicates both positive and negative 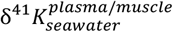 are possible. For example, as Δf_ch_ → 0, 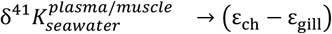 a result which yields 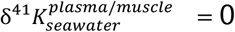 to +1‰, in agreement with measured 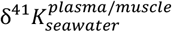 values in euryhaline marine fish. Alternatively, as Δf_ch_ → 1, 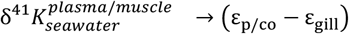 a result which yields 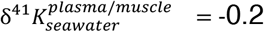 to −1.2‰, in agreement with measured 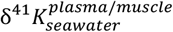 values in stenohaline marine fish.

But is there any independent evidence that K homeostasis in MRCs in stenohaline marine fish operates close to a perfect ‘pump-leak’ (Δf_ch_ ~ 1) mechanism whereas the MRCs of euryhaline marine fish operate much closer to a ‘leak-leak’ or ‘pump-pump’ mechanism (Δf_ch_ ~ 0)? The gill epithelium of stenohaline marine fish is considered to be more permeable than stenohaline freshwater fish due to the presence of tight-junctions connecting gill accessory cells with MRCs (Chasiotis et al., 2012). This increased permeability to NaCl must be balanced by increased NaCl secretion, a process that has been shown to be linked to the activity of NaKATP_ase_ in MRCs; activities of NaKATP_ase_ is higher in marine species than freshwater species and increase when euryhaline species are acclimated to seawater (Epstein et al., 1967; Jampol and Epstein, 1970). As an increase in the activity of NaKATP_ase_ activity will, all else held constant, lower 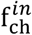 and increase Δf_ch_, one way of interpreting the apparent difference in Δf_ch_ between MRCs of stenohaline and euryhaline marine fish is that the more permeable gills of stenohaline marine fish require greater NaCl secretion and higher activities of NaKATP_ase_ in MRCs (e.g. Figure 3c).

## 5. Conclusions

The results presented here demonstrate that K^+^ homeostasis in biological systems is associated with systematic variability in ^41^K/^39^K ratios and strongly suggests that K^+^ transport through channels and tight-junction proteins is associated with greater fractionation of K isotopes than transport via pumps and co-transporters. Using results from 3 biological systems (algae, mammals, and fish) we developed a simple framework for interpreting differences in measured δ^41^K values between intercellular fluid (ICF) and extracellular fluid (ECF) and between different internal fluid reservoirs (e.g. blood plasma and cerebrospinal fluid). If correct, this framework permits quantification of the relative importance of K^+^ channels and pump/cotransporters in different physiological states – e.g. K depletion/excess - *in situ* (e.g 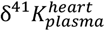) and in some cases *in vivo* (e.g 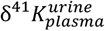). However, with the exception of *C. Reinhardtii* where the observed isotopic difference between media and the whole cell can be attributed to a single transport mechanism, our results do not directly constrain the magnitude of the individual K isotope effects associated with the machinery of K^+^ transport. Quantifying machinery-specific K isotope effects through a combination of laboratory and numerical approaches (14) is therefore a high-priority for future research.

## Data Availability Statement

The data that support the findings of this study are available in the methods and/or supplementary material of this article.

## Author Contributions

JAH, AAM, JHY, DPS, and CS designed the research. JAH, DPS, DH, SK, SG, and CS carried out the experiments and associated analytical measurements. JAH, AAM, JHY, DPS, and CS interpreted the data and wrote the manuscript. All authors were involved in the revising of the manuscript.

## Acknowledgments

Cornelia Spetea acknowledges the sabbatical program at the Faculty of Science, University of Gothenburg, and thanks Martin C. Jonikas for the *Chlamydomonas* experiments performed in his laboratory at Princeton University. John Higgins, Alicia McDonough, and Jang Youn thank the University Kidney Research Organization (UKRO) for financial support. Anne Pearson is thanked for helpful reviews on an earlier version of this manuscript.

---

**Table S1.**
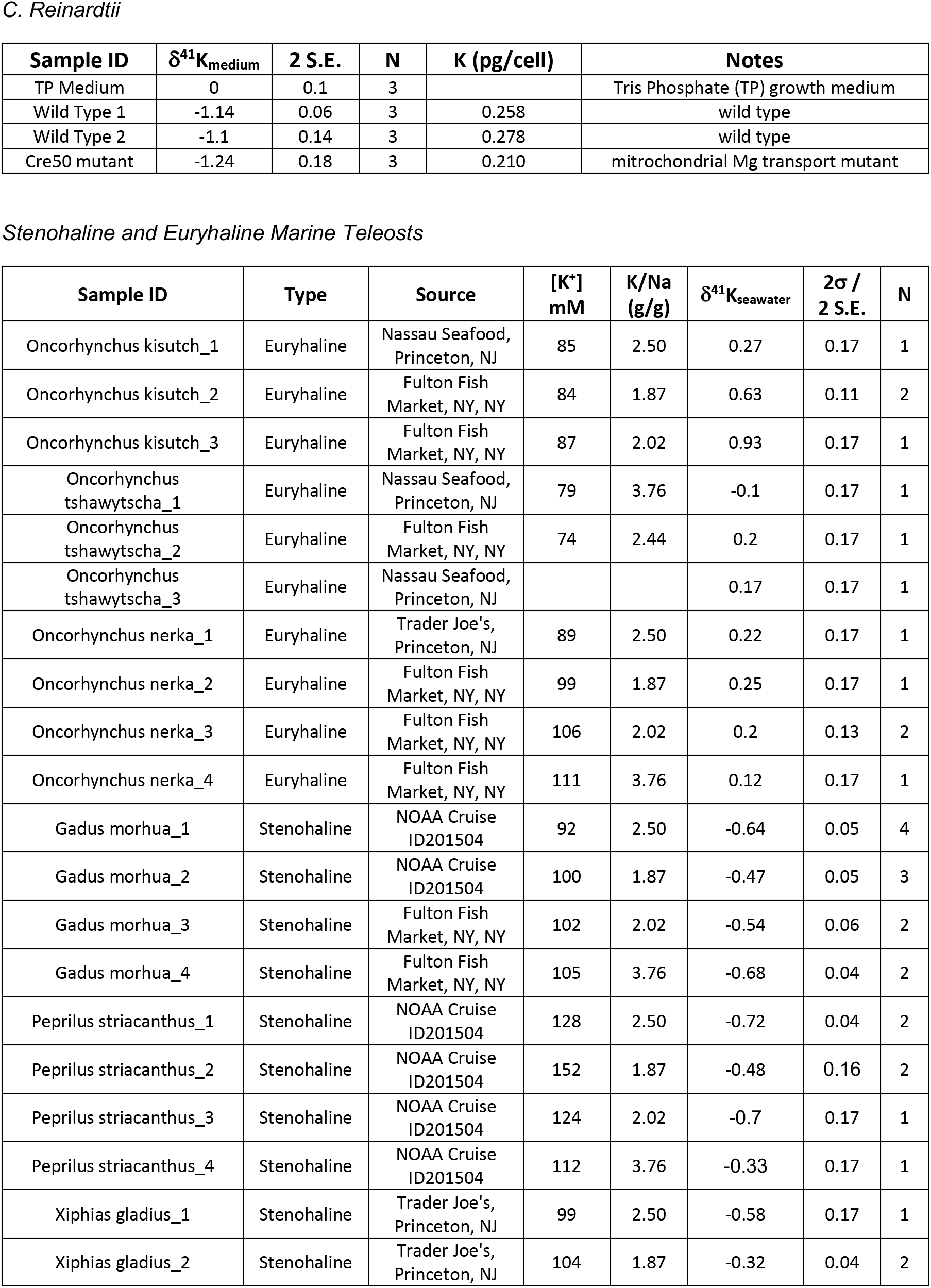
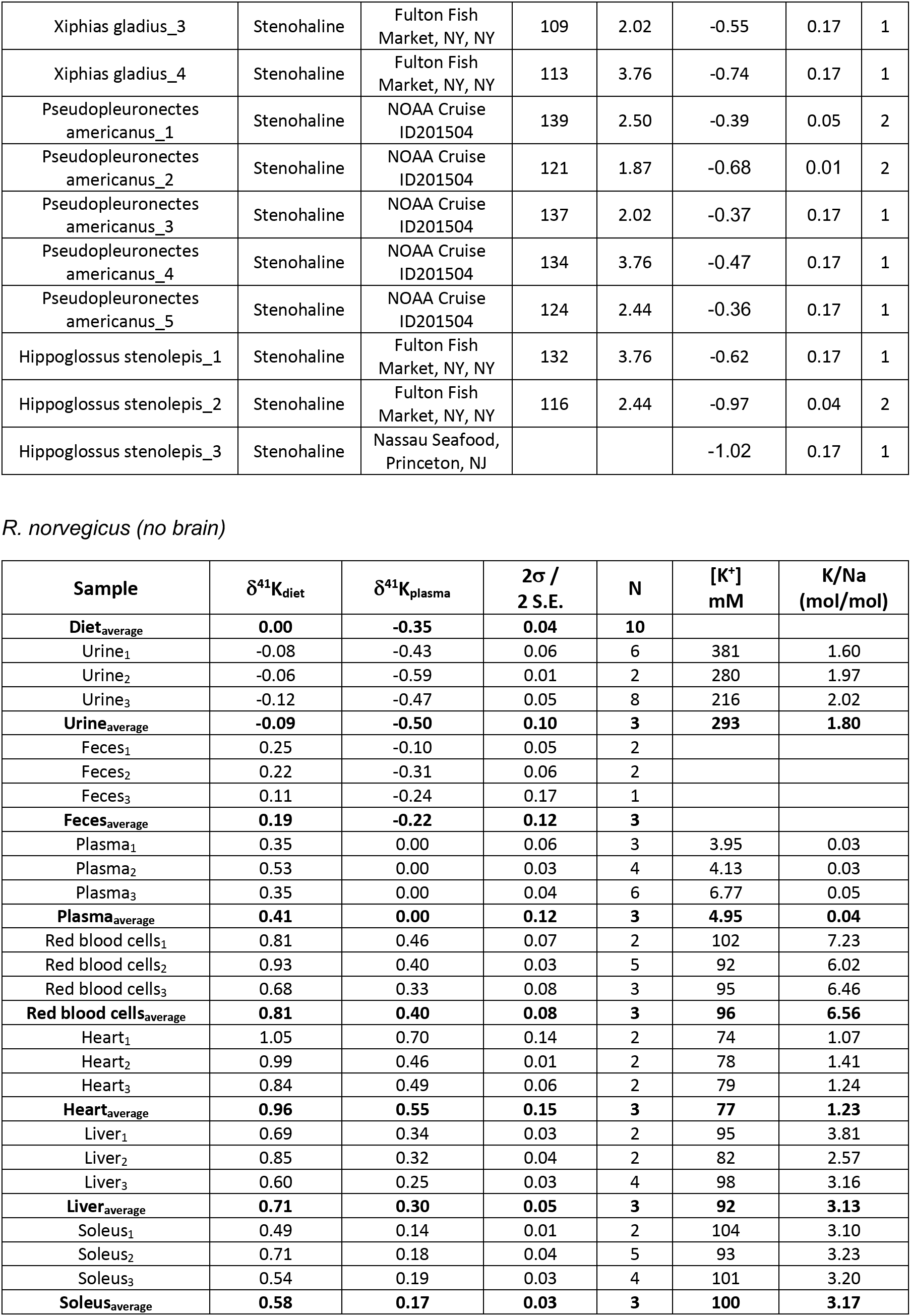
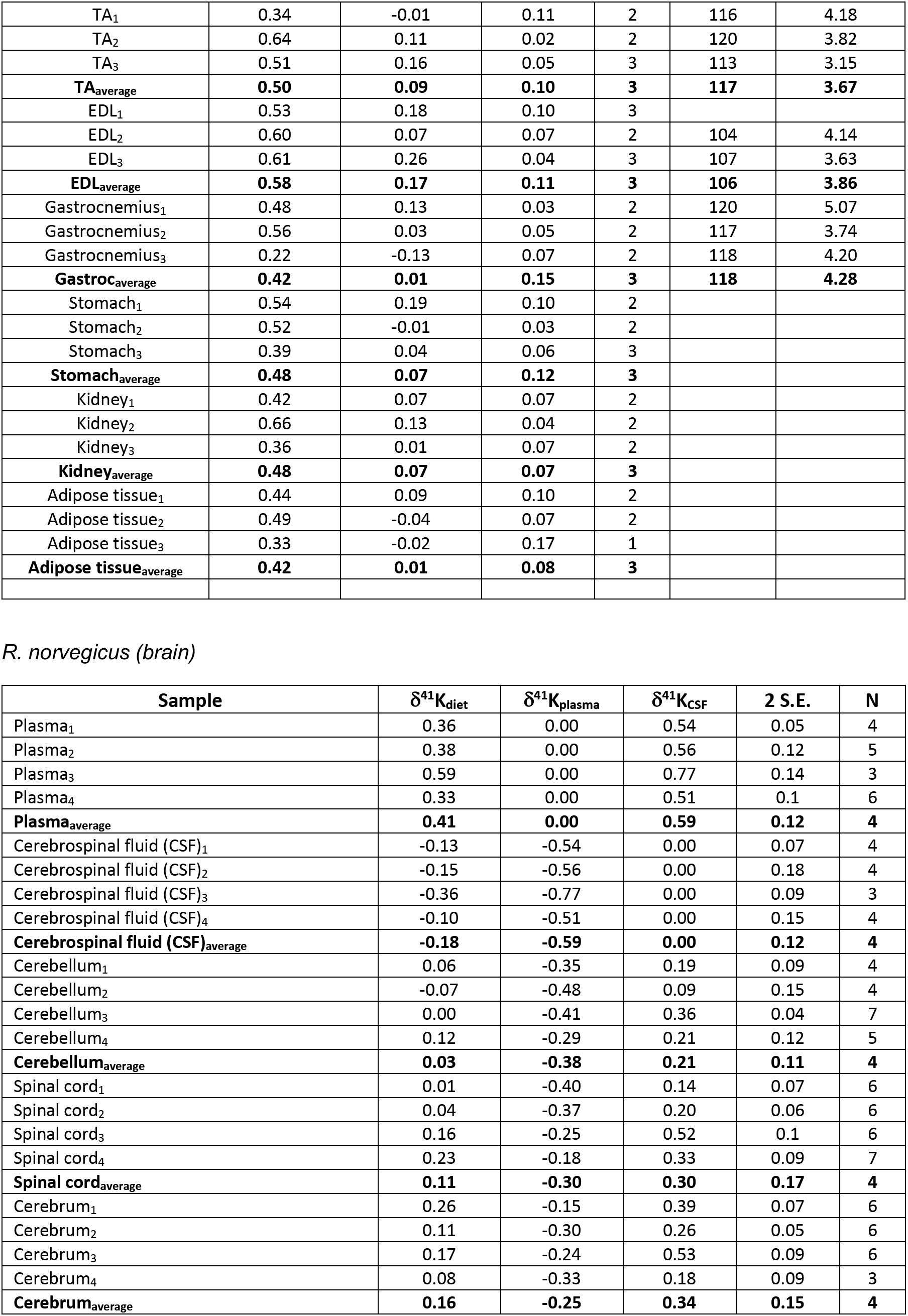

