## Supplemental Table 1 for "Stable Potassium Isotopes (^41^K/^39^K) Track Transcellular and Paracellular Potassium Transport in Biological Systems"

**Table S1**. Measured $\delta^{41}K$ values and other relevant information for all samples in this study.

*C. Reinardtii*

| Sample ID | $\delta^{41}K_{medium}$ | 2 S.E. | N | K (pg/cell) | Notes |
|---|---|---|---|---|---|
| TP Medium | 0 | 0.1 | 3 | | Tris Phosphate (TP) growth medium |
| Wild Type 1 | -1.14 | 0.06 | 3 | 0.258 | wild type |
| Wild Type 2 | -1.1 | 0.14 | 3 | 0.278 | wild type |
| Cre50 mutant | -1.24 | 0.18 | 3 | 0.210 | mitrochondrial Mg transport mutant |

*Stenohaline and Euryhaline Marine Teleosts*

| Sample ID | Type | Source | $[K^+]$ mM | K/Na (g/g) | $\delta^{41}K_{seawater}$ | $2\sigma$ / 2 S.E. | N |
|---|---|---|---|---|---|---|---|
| Oncorhynchus kisutch_1 | Euryhaline | Nassau Seafood, Princeton, NJ | 85 | 2.50 | 0.27 | 0.17 | 1 |
| Oncorhynchus kisutch_2 | Euryhaline | Fulton Fish Market, NY, NY | 84 | 1.87 | 0.63 | 0.11 | 2 |
| Oncorhynchus kisutch_3 | Euryhaline | Fulton Fish Market, NY, NY | 87 | 2.02 | 0.93 | 0.17 | 1 |
| Oncorhynchus tshawytscha_1 | Euryhaline | Nassau Seafood, Princeton, NJ | 79 | 3.76 | -0.1 | 0.17 | 1 |
| Oncorhynchus tshawytscha_2 | Euryhaline | Fulton Fish Market, NY, NY | 74 | 2.44 | 0.2 | 0.17 | 1 |
| Oncorhynchus tshawytscha_3 | Euryhaline | Nassau Seafood, Princeton, NJ | | | 0.17 | 0.17 | 1 |
| Oncorhynchus nerka_1 | Euryhaline | Trader Joe's, Princeton, NJ | 89 | 2.50 | 0.22 | 0.17 | 1 |
| Oncorhynchus nerka_2 | Euryhaline | Fulton Fish Market, NY, NY | 99 | 1.87 | 0.25 | 0.17 | 1 |
| Oncorhynchus nerka_3 | Euryhaline | Fulton Fish Market, NY, NY | 106 | 2.02 | 0.2 | 0.13 | 2 |
| Oncorhynchus nerka_4 | Euryhaline | Fulton Fish Market, NY, NY | 111 | 3.76 | 0.12 | 0.17 | 1 |
| Gadus morhua_1 | Stenohaline | NOAA Cruise ID201504 | 92 | 2.50 | -0.64 | 0.05 | 4 |

| | | | | | | | |
|---|---|---|---|---|---|---|---|
| Gadus morhua_2 | Stenohaline | NOAA Cruise ID201504 | 100 | 1.87 | -0.47 | 0.05 | 3 |
| Gadus morhua_3 | Stenohaline | Fulton Fish Market, NY, NY | 102 | 2.02 | -0.54 | 0.06 | 2 |
| Gadus morhua_4 | Stenohaline | Fulton Fish Market, NY, NY | 105 | 3.76 | -0.68 | 0.04 | 2 |
| Peprilus striacanthus_1 | Stenohaline | NOAA Cruise ID201504 | 128 | 2.50 | -0.72 | 0.04 | 2 |
| Peprilus striacanthus_2 | Stenohaline | NOAA Cruise ID201504 | 152 | 1.87 | -0.48 | 0.16 | 2 |
| Peprilus striacanthus_3 | Stenohaline | NOAA Cruise ID201504 | 124 | 2.02 | -0.7 | 0.17 | 1 |
| Peprilus striacanthus_4 | Stenohaline | NOAA Cruise ID201504 | 112 | 3.76 | -0.33 | 0.17 | 1 |
| Xiphias gladius_1 | Stenohaline | Trader Joe's, Princeton, NJ | 99 | 2.50 | -0.58 | 0.17 | 1 |
| Xiphias gladius_2 | Stenohaline | Trader Joe's, Princeton, NJ | 104 | 1.87 | -0.32 | 0.04 | 2 |
| Xiphias gladius_3 | Stenohaline | Fulton Fish Market, NY, NY | 109 | 2.02 | -0.55 | 0.17 | 1 |
| Xiphias gladius_4 | Stenohaline | Fulton Fish Market, NY, NY | 113 | 3.76 | -0.74 | 0.17 | 1 |
| Pseudopleuronectes americanus_1 | Stenohaline | NOAA Cruise ID201504 | 139 | 2.50 | -0.39 | 0.05 | 2 |
| Pseudopleuronectes americanus_2 | Stenohaline | NOAA Cruise ID201504 | 121 | 1.87 | -0.68 | 0.01 | 2 |
| Pseudopleuronectes americanus_3 | Stenohaline | NOAA Cruise ID201504 | 137 | 2.02 | -0.37 | 0.17 | 1 |
| Pseudopleuronectes americanus_4 | Stenohaline | NOAA Cruise ID201504 | 134 | 3.76 | -0.47 | 0.17 | 1 |
| Pseudopleuronectes americanus_5 | Stenohaline | NOAA Cruise ID201504 | 124 | 2.44 | -0.36 | 0.17 | 1 |
| Hippoglossus stenolepis_1 | Stenohaline | Fulton Fish Market, NY, NY | 132 | 3.76 | -0.62 | 0.17 | 1 |
| Hippoglossus stenolepis_2 | Stenohaline | Fulton Fish Market, NY, NY | 116 | 2.44 | -0.97 | 0.04 | 2 |
| Hippoglossus stenolepis_3 | Stenohaline | Nassau Seafood, Princeton, NJ | | | -1.02 | 0.17 | 1 |

7 *R. norvegicus (no brain)*

| Sample | $\delta^{41}K_{diet}$ | $\delta^{41}K_{plasma}$ | 2σ / 2 S.E. | N | [$K^+$] mM | K/Na (mol/mol) |
|---|---|---|---|---|---|---|
| **Diet$_{average}$** | **0.00** | **-0.35** | **0.04** | **10** | | |
| Urine$_1$ | -0.08 | -0.43 | 0.06 | 6 | 381 | 1.60 |
| Urine$_2$ | -0.06 | -0.59 | 0.01 | 2 | 280 | 1.97 |
| Urine$_3$ | -0.12 | -0.47 | 0.05 | 8 | 216 | 2.02 |
| **Urine$_{average}$** | **-0.09** | **-0.50** | **0.10** | **3** | **293** | **1.80** |
| Feces$_1$ | 0.25 | -0.10 | 0.05 | 2 | | |
| Feces$_2$ | 0.22 | -0.31 | 0.06 | 2 | | |
| Feces$_3$ | 0.11 | -0.24 | 0.17 | 1 | | |
| **Feces$_{average}$** | **0.19** | **-0.22** | **0.12** | **3** | | |
| Plasma$_1$ | 0.35 | 0.00 | 0.06 | 3 | 3.95 | 0.03 |
| Plasma$_2$ | 0.53 | 0.00 | 0.03 | 4 | 4.13 | 0.03 |
| Plasma$_3$ | 0.35 | 0.00 | 0.04 | 6 | 6.77 | 0.05 |
| **Plasma$_{average}$** | **0.41** | **0.00** | **0.12** | **3** | **4.95** | **0.04** |
| Red blood cells$_1$ | 0.81 | 0.46 | 0.07 | 2 | 102 | 7.23 |
| Red blood cells$_2$ | 0.93 | 0.40 | 0.03 | 5 | 92 | 6.02 |
| Red blood cells$_3$ | 0.68 | 0.33 | 0.08 | 3 | 95 | 6.46 |
| **Red blood cells$_{average}$** | **0.81** | **0.40** | **0.08** | **3** | **96** | **6.56** |
| Heart$_1$ | 1.05 | 0.70 | 0.14 | 2 | 74 | 1.07 |
| Heart$_2$ | 0.99 | 0.46 | 0.01 | 2 | 78 | 1.41 |
| Heart$_3$ | 0.84 | 0.49 | 0.06 | 2 | 79 | 1.24 |
| **Heart$_{average}$** | **0.96** | **0.55** | **0.15** | **3** | **77** | **1.23** |
| Liver$_1$ | 0.69 | 0.34 | 0.03 | 2 | 95 | 3.81 |
| Liver$_2$ | 0.85 | 0.32 | 0.04 | 2 | 82 | 2.57 |
| Liver$_3$ | 0.60 | 0.25 | 0.03 | 4 | 98 | 3.16 |
| **Liver$_{average}$** | **0.71** | **0.30** | **0.05** | **3** | **92** | **3.13** |
| Soleus$_1$ | 0.49 | 0.14 | 0.01 | 2 | 104 | 3.10 |

| | | | | | | |
|---|---|---|---|---|---|---|
| Soleus$_2$ | 0.71 | 0.18 | 0.04 | 5 | 93 | 3.23 |
| Soleus$_3$ | 0.54 | 0.19 | 0.03 | 4 | 101 | 3.20 |
| **Soleus$_{average}$** | **0.58** | **0.17** | **0.03** | **3** | **100** | **3.17** |
| TA$_1$ | 0.34 | -0.01 | 0.11 | 2 | 116 | 4.18 |
| TA$_2$ | 0.64 | 0.11 | 0.02 | 2 | 120 | 3.82 |
| TA$_3$ | 0.51 | 0.16 | 0.05 | 3 | 113 | 3.15 |
| **TA$_{average}$** | **0.50** | **0.09** | **0.10** | **3** | **117** | **3.67** |
| EDL$_1$ | 0.53 | 0.18 | 0.10 | 3 | | |
| EDL$_2$ | 0.60 | 0.07 | 0.07 | 2 | 104 | 4.14 |
| EDL$_3$ | 0.61 | 0.26 | 0.04 | 3 | 107 | 3.63 |
| **EDL$_{average}$** | **0.58** | **0.17** | **0.11** | **3** | **106** | **3.86** |
| Gastrocnemius$_1$ | 0.48 | 0.13 | 0.03 | 2 | 120 | 5.07 |
| Gastrocnemius$_2$ | 0.56 | 0.03 | 0.05 | 2 | 117 | 3.74 |
| Gastrocnemius$_3$ | 0.22 | -0.13 | 0.07 | 2 | 118 | 4.20 |
| **Gastroc$_{average}$** | **0.42** | **0.01** | **0.15** | **3** | **118** | **4.28** |
| Stomach$_1$ | 0.54 | 0.19 | 0.10 | 2 | | |
| Stomach$_2$ | 0.52 | -0.01 | 0.03 | 2 | | |
| Stomach$_3$ | 0.39 | 0.04 | 0.06 | 3 | | |
| **Stomach$_{average}$** | **0.48** | **0.07** | **0.12** | **3** | | |
| Kidney$_1$ | 0.42 | 0.07 | 0.07 | 2 | | |
| Kidney$_2$ | 0.66 | 0.13 | 0.04 | 2 | | |
| Kidney$_3$ | 0.36 | 0.01 | 0.07 | 2 | | |
| **Kidney$_{average}$** | **0.48** | **0.07** | **0.07** | **3** | | |
| Adipose tissue$_1$ | 0.44 | 0.09 | 0.10 | 2 | | |
| Adipose tissue$_2$ | 0.49 | -0.04 | 0.07 | 2 | | |
| Adipose tissue$_3$ | 0.33 | -0.02 | 0.17 | 1 | | |
| **Adipose tissue$_{average}$** | **0.42** | **0.01** | **0.08** | **3** | | |

*R. norvegicus (brain)*

| Sample | $\delta^{41}K_{diet}$ | $\delta^{41}K_{plasma}$ | $\delta^{41}K_{CSF}$ | 2 S.E. | N |
|---|---|---|---|---|---|
| Plasma$_1$ | 0.36 | 0.00 | 0.54 | 0.05 | 4 |
| Plasma$_2$ | 0.38 | 0.00 | 0.56 | 0.12 | 5 |
| Plasma$_3$ | 0.59 | 0.00 | 0.77 | 0.14 | 3 |
| Plasma$_4$ | 0.33 | 0.00 | 0.51 | 0.1 | 6 |
| **Plasma$_{average}$** | **0.41** | **0.00** | **0.59** | **0.12** | **4** |
| Cerebrospinal fluid (CSF)$_1$ | -0.13 | -0.54 | 0.00 | 0.07 | 4 |
| Cerebrospinal fluid (CSF)$_2$ | -0.15 | -0.56 | 0.00 | 0.18 | 4 |
| Cerebrospinal fluid (CSF)$_3$ | -0.36 | -0.77 | 0.00 | 0.09 | 3 |
| Cerebrospinal fluid (CSF)$_4$ | -0.10 | -0.51 | 0.00 | 0.15 | 4 |
| **Cerebrospinal fluid (CSF)$_{average}$** | **-0.18** | **-0.59** | **0.00** | **0.12** | **4** |
| Cerebellum$_1$ | 0.06 | -0.35 | 0.19 | 0.09 | 4 |
| Cerebellum$_2$ | -0.07 | -0.48 | 0.09 | 0.15 | 4 |
| Cerebellum$_3$ | 0.00 | -0.41 | 0.36 | 0.04 | 7 |
| Cerebellum$_4$ | 0.12 | -0.29 | 0.21 | 0.12 | 5 |
| **Cerebellum$_{average}$** | **0.03** | **-0.38** | **0.21** | **0.11** | **4** |
| Spinal cord$_1$ | 0.01 | -0.40 | 0.14 | 0.07 | 6 |
| Spinal cord$_2$ | 0.04 | -0.37 | 0.20 | 0.06 | 6 |
| Spinal cord$_3$ | 0.16 | -0.25 | 0.52 | 0.1 | 6 |
| Spinal cord$_4$ | 0.23 | -0.18 | 0.33 | 0.09 | 7 |
| **Spinal cord$_{average}$** | **0.11** | **-0.30** | **0.30** | **0.17** | **4** |
| Cerebrum$_1$ | 0.26 | -0.15 | 0.39 | 0.07 | 6 |
| Cerebrum$_2$ | 0.11 | -0.30 | 0.26 | 0.05 | 6 |
| Cerebrum$_3$ | 0.17 | -0.24 | 0.53 | 0.09 | 6 |
| Cerebrum$_4$ | 0.08 | -0.33 | 0.18 | 0.09 | 3 |
| **Cerebrum$_{average}$** | **0.16** | **-0.25** | **0.34** | **0.15** | **4** |

12
